# Traumatic brain injury alters dendritic cell differentiation and distribution in lymphoid and non-lymphoid organs

**DOI:** 10.1101/2021.12.28.474349

**Authors:** Orest Tsymbalyuk, Volodymyr Gerzanich, J Marc Simard, Chozha Vendan Rathinam

## Abstract

Pathophysiological consequences of traumatic brain injury (TBI) mediated secondary injury remain incompletely understood. In particular, the impact of TBI on the differentiation and maintenance of dendritic cells (DCs), remains completely unknown. Here, we report that DC-differentiation, maintenance and functions are altered at both early and late phases of TBI. Our studies identify that; 1. frequencies and absolute numbers of DCs in the spleen and BM are altered at both acute and late phases of TBI; 2. surface expression of key molecules involved in antigen presentation of DCs were affected both at early and late phases of TBI; 3. distribution and functions of tissue-specific DC subsets of both circulatory and lymphatic systems were imbalanced following TBI; 4. early differentiation program of DCs, especially the commitment of hematopoietic stem cells to common DC progenitors, were deregulated after TBI; and 5. intracellular ROS levels were reduced in DC progenitors and differentiated DCs at both early and late phases of TBI. Our data demonstrate, for the first time, that TBI affects the distribution pattern of DCs and induces an imbalance among DC subsets in both lymphoid and non-lymphoid organs. In addition, the current study demonstrates that TBI results in reduced levels of ROS in DCs at both early and late phases of TBI, which may explain altered DC differentiation paradigm following TBI. A deeper understanding on the molecular mechanisms that contribute to DC defects following TBI would be essential and beneficial in treating infections in patients with acute central nervous system (CNS) injuries.

## Background

Traumatic brain injury (TBI)-mediated disabilities and/or deaths pose major threats to global health [1, 2]. TBI affects ∼ 1.7 million people each year and contributes to a third of all injury related deaths within the United States [1, 3-6]. Even though cellular and molecular mediators that cause severe trauma within the central nervous system (CNS) following TBI have been widely recognized in the recent years, the systemic impact of TBI remains incompletely understood. In particular, knowledge on the cascade of cellular and molecular events leading to TBI-induced “secondary injuries” [7] and its pathophysiological consequences remain elusive.

Dendritic cells (DCs) are the most potent antigen presenting cells and function as the sentinels of the immune system. DCs initiate and shape both innate and adaptive immune responses[8, 9]. In particular, DCs are essential for the initiation of protective T- and B-cell responses and, thus constitute a frontline defense against invading pathogens[10-12]. Indeed, impaired DC differentiation and functions ultimately results in severe autoimmune deficiencies and enhanced susceptibility to viral, bacterial and fungal infections in mice and humans[13-15]. DCs are present in most tissues of the body and can be broadly divided into 3 major groups; 1. conventional or classical DCs (cDCs), which are further subdivided into cDC1 and cDC2 subsets, 2. plasmacytoid DCs (pDCs) and 3. monocyte-derived DCs (moDCs)[11, 16, 17]. cDCs patrol the local environment, actively engulf and sample foreign antigens, migrate to the T-cell zones of the draining lymph nodes, and present antigenic peptides to antigen inexperienced/naïve T cells. pDCs are, particularly, essential for the production of type I interferons and the establishment of anti-viral immunity. On the other hand, moDCs are routinely generated *in vitro* from monocytes in presence of granulocyte-macrophage colony stimulating factor (GM-CSF, CSF-2) and interleukin4 (IL4) and are often utilized under clinical settings to perform immunotherapies against cancer[17].

Earlier studies unequivocally established the impact of TBI on peripheral immune cells, including both innate (granulocytes and macrophages) and adaptive (T-, B-, and NK-cells) immune system[18-21]. A series of recent studies highlighted the significance of DC-mediated functions during CNS injuries, including TBI[22], middle cerebral artery occlusion and cerebral ischemia[23]. However, physiological consequences of TBI on the development of DCs remain totally unknown. Our previous studies[24-27], as well as of others, established that sustained and chronic inflammation impairs early differentiation pathways in HSCs. In particular, our studies indicated that the Flt3^+^ hematopoietic progenitors are more sensitive to acute and chronic inflammation. Based on the key roles of Flt3/Flt3L signaling axis in the differentiation and maintenance of DCs, we hypothesized that TBI-induced inflammation affects differentiation and maintenance of peripheral DCs.

In the present study, we induced experimental TBI in mice and studied its impact on DC maintenance. Our data identified that TBI affects the distribution pattern of DCs and induces an imbalance among DC subsets in both lymphoid and non-lymphoid organs. In addition, the current study demonstrates that TBI results in reduced levels of reactive oxygen species (ROS) in DCs at both early and late phases of TBI, which may explain altered DC differentiation paradigm following TBI.

## Materials and Methods

### Mice and controlled cortical impact injury

Mice were analyzed between 4-12 weeks after birth, unless otherwise specified. C57BL/6 mice were purchased from the Jackson Laboratory. 8-12-week mice were used in this study. After being anesthetized with isoflurane, the subject mice received controlled cortical impact (CCI) using a custom microprocessor-controlled and compressed air driven pneumatic impactor. Briefly, a 10-mm midline incision was made over the skull, the skin and fascia were retracted, and a 4-mm craniotomy was made on the central aspect of the left parietal bone of mice under surgical anesthesia. A moderate injury was induced by a 3.5-mm diameter tip with impact velocity of 6 m/s and a deformation depth of 2 mm. In sham mice, same procedure was performed except for the impact. The number of mice in each study is indicated in the figure legends. All surgical procedures and animal experiments were performed under protocols approved by the University of Maryland School of Medicine Institutional Animal Care and Use Committee (IACUC). The surgical procedures were performed by the same investigator using the same equipment.

### Cell preparation

Spleen, lymph nodes (LNs), liver, lungs were digested with Collagenase-D (0.7 mg/mL) and DNase I (100U/ mL) in RPMI for 45 mins at 37° C. For skin DC isolation, mouse ears were cut into pieces, digested in the presence of 250 ug/mL liberase + DNase I (100U/ mL) in HBSS + 5% FCS for 90 mins at 37° C. Cells were vortexed vigorously and passed through a 19-gauge needle to obtain a single cell suspension. Bone marrow cells were isolated from the tibias and femurs by inserting a 23-gauge needle/ 1 mL syringe to the bone cavities and flushed with PBS 2%FCS until the bones become pale. Single cell suspensions were made through rigorous pipetting.

Red blood cells were lysed with Ammonium chloride (Stem Cell Technology) and subsequently filtered using a 70 μM nylon mesh. Bone marrow cells were then counted with a hemacytometer and trypan blue (Amresco) negative cells were counted as live cells.

### Flow cytometry

Cells were analyzed by flow cytometry with Attune Nxt (Thermofisher) and FlowJo software (Tree Star). The following monoclonal antibodies were used: anti-CD34 (RAM34), anti-CD48 (HM48-1), anti-CD117 (2B8), anti-Flt3 (A2F10.1), anti-Sca-1 (D7), anti-B220 (RA3-6B2), anti-CD19 (1D3), anti-CD3 (145-2C11), anti-CD4 (GK1.5), anti-CD8 (53-6.7), anti-CD11b (M1/70), anti– Gr-1 (RB6-8C5), anti-Ter119 (TER119), anti-CD11c (N418), anti-CD8(53-6.7), anti-PDCA1 (129C1), anti-CD24 (M1/69), anti-SIRPa (P84), anti-CD103 (2E7), anti-CD45 (S18009F), anti-CD207 (4C7), anti-CD115 (AFS98), anti-LY6C (HK1.4), anti-SiglecH (551), anti-CLEC9A (7H11), anti-CD80 (16-10A1), anti-CD86 (GL-1), anti-CD40(3/23), anti-H2K^b^ (AF6-88.5), anti-IA/IE (M5/114.15.2) and anti-BrdU (3D4) from Biolegend. Cells incubated with biotinylated monoclonal antibodies were incubated with fluorochrome-conjugated streptavidin–peridinin chlorophyll protein–cyanine 5.5 (551419; BD), streptavidin-allophycocyanin-Cy7 (554063; BD), streptavidin-super bright 650 (Biolegend). In all the FACS plots, indicated are the percentages (%) of the gated fraction. Data were acquired on a Attune Nxt Acoustic focusing cytometer using Attune software (life technologies) and analyzed using FlowJo (Treestar Inc).

### Measuring ROS levels

Single cell suspensions were stained with cell surface markers and then incubated with 2mM CM-H_2_DCFDA (Life Technologies C6827) in pre-warmed HBSS at 37C for 15 min. The cells were washed in ice-cold PBS, pelleted and resuspended in ice-cold PBS 2 % FCS.

### Statistics

Data represent mean and s.e.m. Significance was evaluated between 2 individual samples using Student unpaired *t-*tests. Comparisons within each surgery group were analyzed using two-way ANOVA with multiple comparisons test. For non-parametric data, Mann Whitney test was used (*P < 0.05, **P<0.01, *** < 0.001, **** < 0.0001).

## Results

### Augmented DC pool size in the spleen and bone marrow at the acute phase of TBI

To study if TBI causes alterations in DC compartment, we enumerated the frequencies of DCs in the spleen and BM. Flow cytometry-based immunophenotyping studies indicated reduced relative frequencies, but increased absolute numbers, of CD11c^+^CII^+^ cDCs in the spleen after 3 days (d3) of TBI (**Fig. 1A-B**). Further fractionation of CD11c^+^CII^+^ DCs into CD8^+^CD11b^-^ (cDC1) and CD8^-^CD11b^+^ (cDC2) subsets[11, 16, 17] subsets revealed; normal relative numbers, but reduced absolute numbers of cDC1 after 1 day (d1) of TBI (**Fig. 1C-D**); an increase in relative frequencies, normal total frequencies and increased absolute numbers of cDC1 subset after d3 of TBI (**Fig. 1C-D**); reduced overall frequencies, but increased absolute numbers of cDC2 after d3 of TBI (**Fig. 1C, E**); and normal relative frequencies, reduced overall frequencies and increased absolute numbers of CD8-CD11b-CD11c^+^CII^+^ immature cDCs in the spleen after d3 of TBI (**Fig. 1C-F**). Further analysis indicated reduced relative numbers, but increased absolute numbers, of PDCA1^+^CD11c^int^ pDCs (**Fig. 1G-H**) and PDCA1^-^ CD11c^high^ cDCs (**Fig. 1G, I**) in the spleen after d3 of TBI. Interestingly, normal relative frequencies, but reduced absolute numbers, of PDCA1^-^CD11c^high^ cDCs was observed in the spleen after d1 of TBI (**Fig. 1G, I**). Next, we determined the frequencies of DCs in the BM and analysis revealed an increase in both relative frequencies and absolute numbers of PDCA1^+^CD11c^int^ pDCs in the BM after d1 and d3 of TBI (**Fig. 1J-K**). On the other hand, frequencies of CD11c^+^CII^+^ cDCs were reduced, even though their absolute numbers were normal, in the BM after d3 of TBI (**Fig. 1L-M**). Finally, absolute numbers of CD11c^+^CII^+^ cDCs were increased, while the relative numbers were normal, in the BM after d1 of TBI (**Fig. 1L-M**). Overall, these data indicate that the frequencies and absolute numbers of DCs in the spleen and BM are altered at the acute phase of TBI.

**Figure 1.**
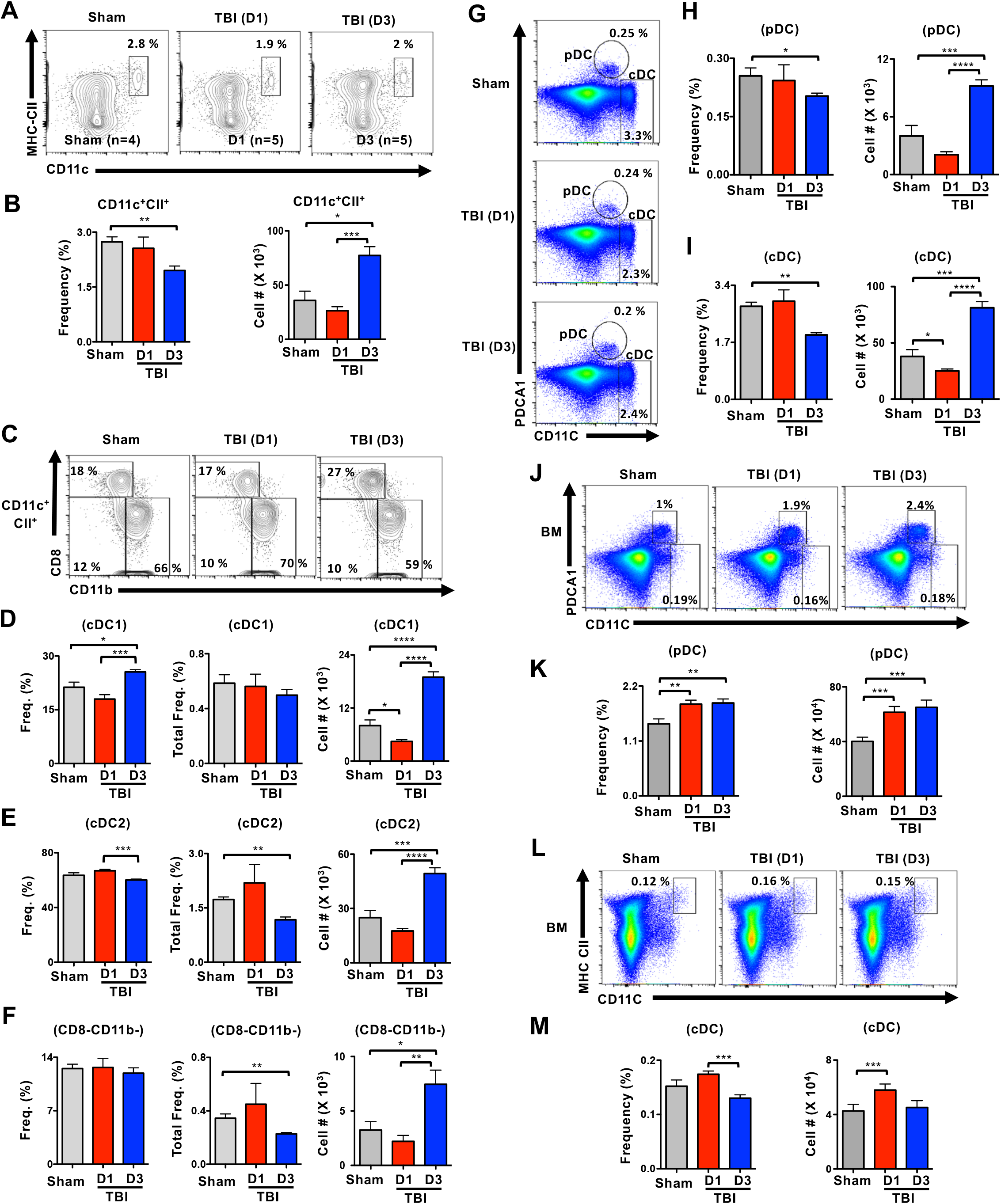
TBI leads to altered numbers and distribution of DC subsets at the early phase. **A**. FACS plots indicating frequencies of CD11c^+^MHC-Class II^high^ cDCs in the spleen of Sham, d1-TBI and d3-TBI mice. Data are representative of two independent experiments. **B**. Cumulative frequencies (**left**) and absolute numbers (**right**) of cDCs in the spleen of Sham (n=4), d1-TBI (n=5) and d3-TBI (n=5) mice. Data are representative of two independent experiments. **C**. FACS plots indicating frequencies of CD8^+^CD11b^-^cDC1, CD8^-^CD11b^+^cDC2 and CD8^-^ CD11b^-^ immature cDCs within pre-gated CD11c^+^MHC-Class II^high^ cDCs in the spleen of Sham (n=4), d1-TBI (n=5) and d3-TBI (n=5) mice. Data are representative of two independent experiments. **D-F**. Relative frequencies (**left**), overall frequencies (**middle**) and absolute numbers (**right**) of cDC1 (**D)**, cDC2 (**E)** and immature cDCs (**F)** subsets in the spleen of Sham (n=4), d1-TBI (n=5) and d3-TBI (n=5) mice. Data are representative of two independent experiments. **G**. FACS plots indicating frequencies of PDCA1^+^CD11c^int^ pDCs and PDCA1^-^CD11c^high^ cDCs in the spleen of Sham (n=4), d1-TBI (n=5) and d3-TBI (n=5) mice. Data are representative of two independent experiments. **H & I**. Overall frequencies (**left**) and absolute numbers (**right**) of pDCs (**H)** and cDCs (**I)** in the spleen of Sham (n=4), d1-TBI (n=5) and d3-TBI (n=5) mice. Data are representative of two independent experiments. **J**. FACS plots indicating frequencies of PDCA1^+^CD11c^int^ pDCs and PDCA1^-^CD11c^high^ cDCs in the BM of Sham (n=4), d1-TBI (n=5) and d3-TBI (n=5) mice. Data are representative of two independent experiments. **K**. Overall frequencies (**left**) and absolute numbers (**right**) of pDCs in the BM of Sham (n=4), d1-TBI (n=5) and d3-TBI (n=5) mice. Data are representative of two independent experiments. **L**. FACS plots indicating frequencies of CD11c^+^MHC-ClassII^high^ cDCs in the BM of Sham (n=4), d1-TBI (n=5) and d3-TBI (n=5) mice. Data are representative of two independent experiments. **M**. Overall frequencies (**left**) and absolute numbers (**right**) of cDCs in the BM of Sham (n=4), d1-TBI (n=5) and d3-TBI (n=5) mice. Data are representative of two independent experiments. All data represent mean ± SEM. Mann-Whitney non-parametric tests were used to assess statistical significance (*P < 0.05, **P<0.01, *** P< 0.001, **** P< 0.0001).

### Altered DC subsets in the spleen and BM at the late phase of TBI

To investigate the impact of TBI on DC subsets at the later phases, we performed immunophenotyping studies after 7 days (d7) of TBI. Our studies indicated normal frequencies, but increased absolute numbers, of CD11c^+^CII^+^ cDCs in the spleen of d7 TBI mice (**Fig. 2A**). Further analysis of splenic cDC fraction of d7 TBI mice indicated; increased frequencies and absolute numbers of cDC1 subset (**Fig. 2B**); reduced frequencies, but normal absolute numbers, of cDC2 subset (**Fig. 2C**); and reduced frequencies, but normal absolute numbers, of immature cDC subset (**Fig. 2D**). Analysis of pDC compartment indicated increased frequencies and absolute numbers in spleen of d7 TBI mice (**Fig. 2E**). Consistent with data shown in **Fig. 2A**, frequencies of PDCA1^-^CD11c^high^ were normal, but their absolute numbers were augmented in the spleen of d7 TBI mice (**Fig. 2F**).

**Figure 2.**
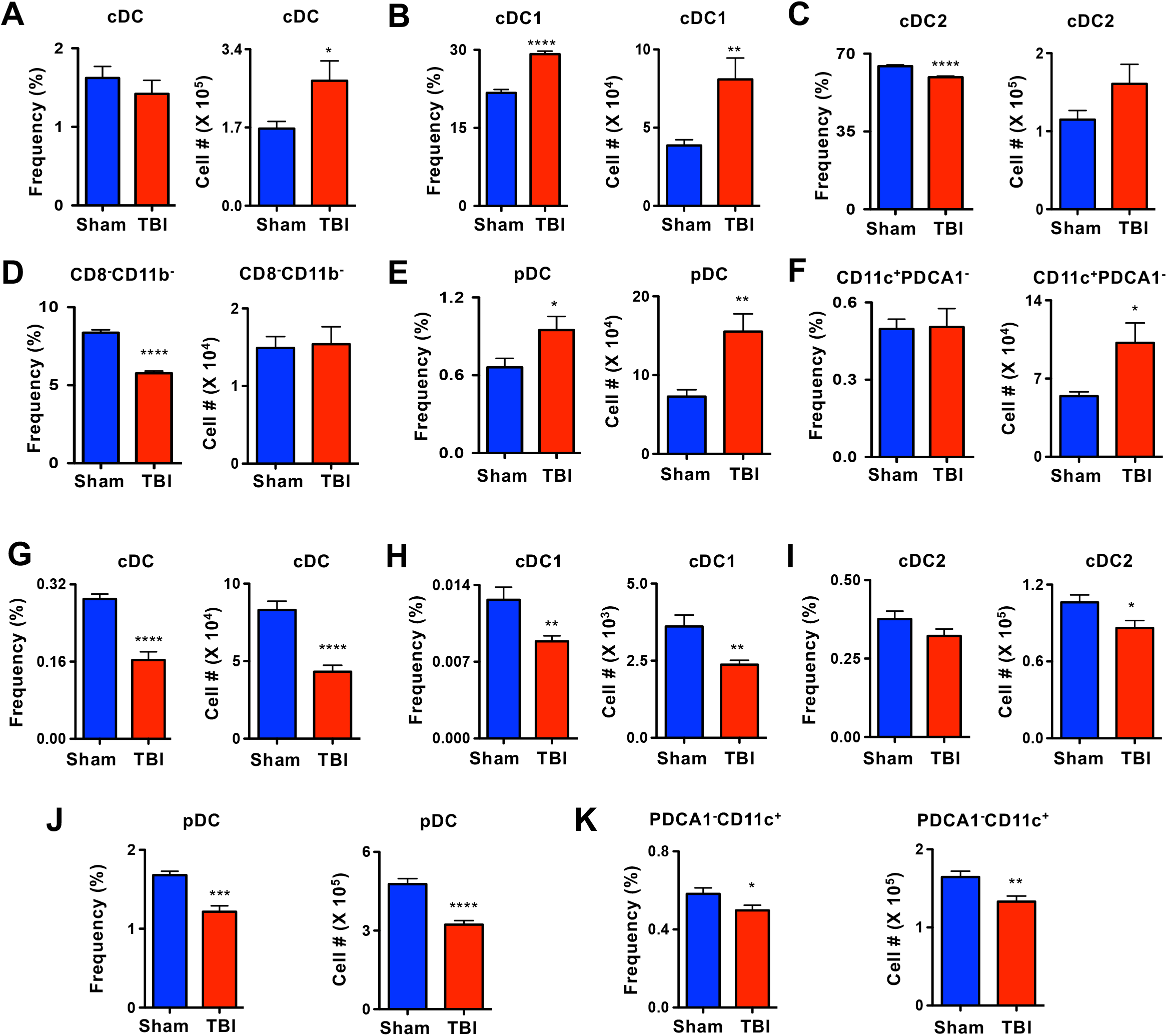
TBI causes changes in DC pool and functions at the late phase. **A**. Relative frequencies (**left**) and absolute numbers (**right**) of cDCs in the spleen of Sham (n=4) and d7-TBI (n=5) mice. Data are representative of two independent experiments. **B**. Relative frequencies (**left**) and absolute numbers (**right**) of cDC1 subset in the spleen of Sham (n=4) and d7-TBI (n=5) mice. Data are representative of two independent experiments. **C**. Relative frequencies (**left**) and absolute numbers (**right**) of cDC2 subset in the spleen of Sham (n=4) and d7-TBI (n=5) mice. Data are representative of two independent experiments. **D**. Relative frequencies (**left**) and absolute numbers (**right**) of immature cDC subset in the spleen of Sham (n=4) and d7-TBI (n=5) mice. Data are representative of two independent experiments. **E**. Relative frequencies (**left**) and absolute numbers (**right**) of pDC subset in the spleen of Sham (n=4) and d7-TBI (n=5) mice. Data are representative of two independent experiments. **F**. Relative frequencies (**left**) and absolute numbers (**right**) of PDCA1^-^CD11c^high^ cDC subset in the spleen of Sham (n=4) and d7-TBI (n=5) mice. Data are representative of two independent experiments. **G**. Relative frequencies (**left**) and absolute numbers (**right**) of cDCs in the BM of Sham (n=4) and d7-TBI (n=5) mice. Data are representative of two independent experiments. **H**. Relative frequencies (**left**) and absolute numbers (**right**) of cDC1 subset in the BM of Sham (n=4) and d7-TBI (n=5) mice. Data are representative of two independent experiments. **I**. Relative frequencies (**left**) and absolute numbers (**right**) of cDC2 subset in the BM of Sham (n=4) and d7-TBI (n=5) mice. Data are representative of two independent experiments. **J**. Relative frequencies (**left**) and absolute numbers (**right**) of pDC subset in the BM of Sham (n=4) and d7-TBI (n=5) mice. Data are representative of two independent experiments. **K**. Relative frequencies (**left**) and absolute numbers (**right**) of PDCA1^-^CD11c^high^ cDC subset in the BM of Sham (n=4) and d7-TBI (n=5) mice. Data are representative of two independent experiments. All data represent mean ± SEM. Two-tailed student’s t tests were used to assess statistical significance (*P < 0.05, **P<0.01, *** P< 0.001, **** P< 0.0001).

On the other hand, analysis of DC subsets in the BM of d7 TBI mice revealed; a remarkable decrease in both frequencies and absolute numbers of CD11c^+^CII^+^ cDCs (**Fig. 2G**); reduced frequencies and absolute numbers of cDC1 (**Fig. 2H**); non-significant reduction in relative frequencies, but a reduction in absolute numbers, of cDC2 (**Fig. 2I**); decreased relative frequencies and absolute numbers of pDCs (**Fig. 2J**); and reduced frequencies and absolute numbers of PDCA1^-^CD11c^high^ cDCs (**Fig. 2K**). In essence, these data specify that later phases of TBI induces an imbalance in the distribution of cDC and pDC subsets in the spleen and BM.

### TBI alters expression levels of surface markers involved in DC functions

To test if expression of surface antigens essential for DC functions is affected by TBI, we performed detailed immunophenotyping studies. Analysis indicated CD80 expression was normal in total DCs, cDC1 and cDC2 subsets of the spleen after d1 and d3 of TBI (**Fig. 3A**). However, CD80 expression was modestly reduced in splenic pDCs after d3, when compared with d1, of TBI (**Fig. 3A**). Expression of CD86 was increased on total DCs and cDC2 subset on d1, when compared with that of d3 after TBI (**Fig. 3B**). Interestingly, CD86 expression levels in cDC1 subset were reduced on d3 after TBI and in pDCs remain unchanged after TBI (**Fig. 3B**). More importantly, expression levels of MHC class I (**Fig. 3C**) and MHC class II (**Fig. 3D**) were normal in total DCs, cDC1, cDC2 and pDCs of spleen after d1 and d3 of TBI.

**Figure 3.**
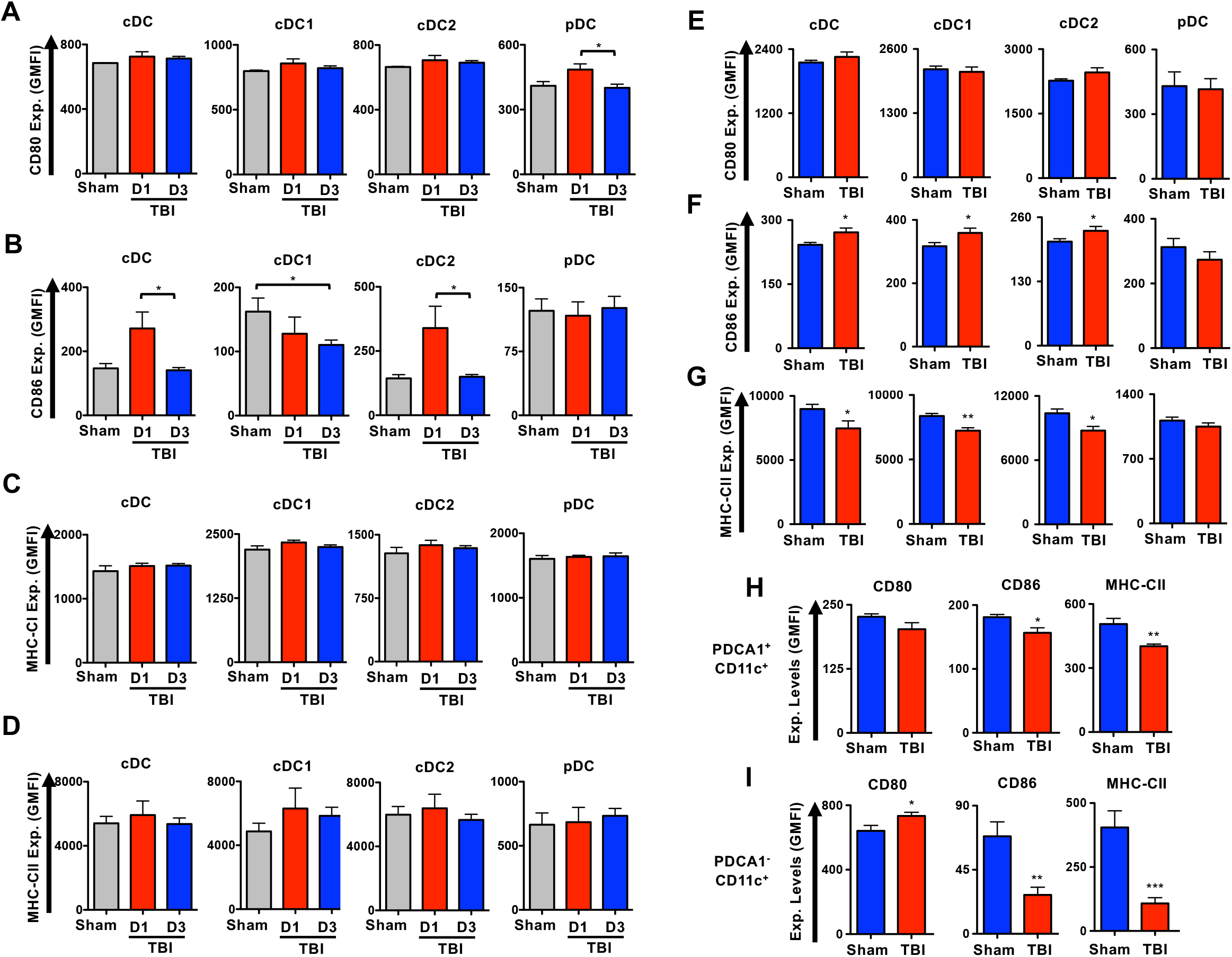
TBI alters expression of expression levels of surface molecules involved in DC functions. **A-D**. Surface expression levels of CD80 (**A**), CD86 (**B**), MHC-CI (**C**) and MHC-CII (**D**) in total cDC, cDC1, cDC2 and pDC subsets of spleen from Sham, d1- and d3-TBI mice. Shown were Geo-Mean Fluorescence Intensities (GMFI). **E-G**. Surface expression levels of CD80 (**E**), CD86 (**F**) and MHC-CII (**G**) in total cDC, cDC1, cDC2 and pDC subsets of spleen from Sham and d7-TBI mice. Shown were GMFI. **H & I**. Surface expression levels of CD80, CD86 and MHC-CII in PDCA1^+^CD11c^+^ pDC (**H**) and PDCA1^-^CD11c^+^ cDC (**I**) subsets of the BM from Sham, d1- and d3-TBI mice. Shown were GMFI. All data represent mean ± SEM. Mann-Whitney non-parametric tests and two-tailed student’s t tests were used to assess statistical significance (*P < 0.05, **P<0.01, *** P< 0.001, **** P< 0.0001).

Next, we assessed if DC associated surface markers are altered at the late phase of TBI. Analysis on expression levels of surface antigen in splenic DC subsets of d7 TBI mice revealed; normal expression levels of CD80 in cDC, cDC1, cDC2 and pDC subsets (**Fig. 3E**); increased expression levels of CD86 in total cDC, cDC1 and cDC2 subsets, whereas normal levels of CD86 in pDCs (**Fig. 3F**); and reduced MHC-CII expression in total cDC, cDC1 and cDC2 subsets, whereas normal levels of MHC-CII in pDCs (**Fig. 3G**). Immunophenotyping studies on BM DC subsets of d7 TBI mice indicated decreased MHC-CII and CD86 expression, whereas normal CD80 expression, in pDC subset (**Fig. 3H**) and striking reduction of MHC-CII and CD86 expression, whereas increased levels of CD80 expression, in PDCA1^-^ CD11C^+^ cDC fraction (**Fig. 3I**). Overall, these data indicate that TBI affects surface expression of molecules involved in antigen presentation of DCs at both early and late phases of injury.

### TBI causes an imbalance in DCs subsets within the circulatory and lymphatic systems

Under steady state conditions DCs are constantly circulated throughout the body to perform immune surveillance and distributed to different organs through sophisticated networks of the blood and lymphatic systems[8-12]. To study if TBI has an impact on circulating DCs, we determined their frequencies in the blood and lymph nodes. Analysis of peripheral blood indicated normal frequencies of total cDCs (**Fig. 4A**). However, within cDCs, a remarkable increase of cDC1 and a decrease of cDC2 subsets were observed in d7 TBI mice (**Fig. 4A**). Interestingly, expression levels of MHC-CII were increased in total cDCs of blood from d7 TBI mice and further analysis indicated that this increase was specific to the cDC1, but not to the cDC2, compartment (**Fig. 4B**). Next, we investigated the frequencies of DCs in the skin draining/peripheral lymph nodes (pLN). Analysis indicated normal frequencies of total cDCs and pDCs in the pLNs from d7 TBI mice (**Fig. 4C**). However, determination of cDC subsets revealed a remarkable decrease of cDC1 and increase of cDC2 subsets in the pLNs of d7 TBI mice (**Fig. 4C**). Interestingly, immunophenotyping studies on pLN DCs from d7 TBI mice indicated; a striking decrease of MHC-CII expression in all DC subsets, including total cDC, cDC1, cDC2 and pDC subsets (**Fig. 4D, left**); normal expression levels of CD80 in total cDCs and pDCs. However, within cDCs, CD80 was decreased in cDC1 and increased in cDC2 subsets (**Fig. 4D, middle**); and CD86 expression was remarkably reduced in total cDC, cDC1 and pDC subsets, even though CD86 expression was normal in cDC2 subset (**Fig. 4D, right**). Finally, we analyzed the DC compartments in the mesenteric lymph nodes (mLNs) and the data indicated that the frequencies of total cDCs were decreased in d7 TBI mice (**Fig. 4E**). However, within cDCs, there was a relative increase of cDC1 subset and a decrease in cDC2 subset in the mLNs of d7 TBI mice (**Fig. 4E**). Immunophenotyping studies on mLN cDCs from d7 TBI mice suggested; normal expression levels of MHC-CII in total cDC, cDC1 and cDC2 subsets (**Fig. 4F, left**); increased expression levels of CD80 expression in cDC, cDC1 and cDC2 subsets (**Fig. 4F, middle**); and remarkable upregulation of CD86 in cDC, cDC1 and cDC2 subsets (**Fig. 4F, right**). Taken together these data demonstrate that TBI affects distribution and functions of specific DC subsets of both circulatory and lymphatic systems.

**Figure 4.**
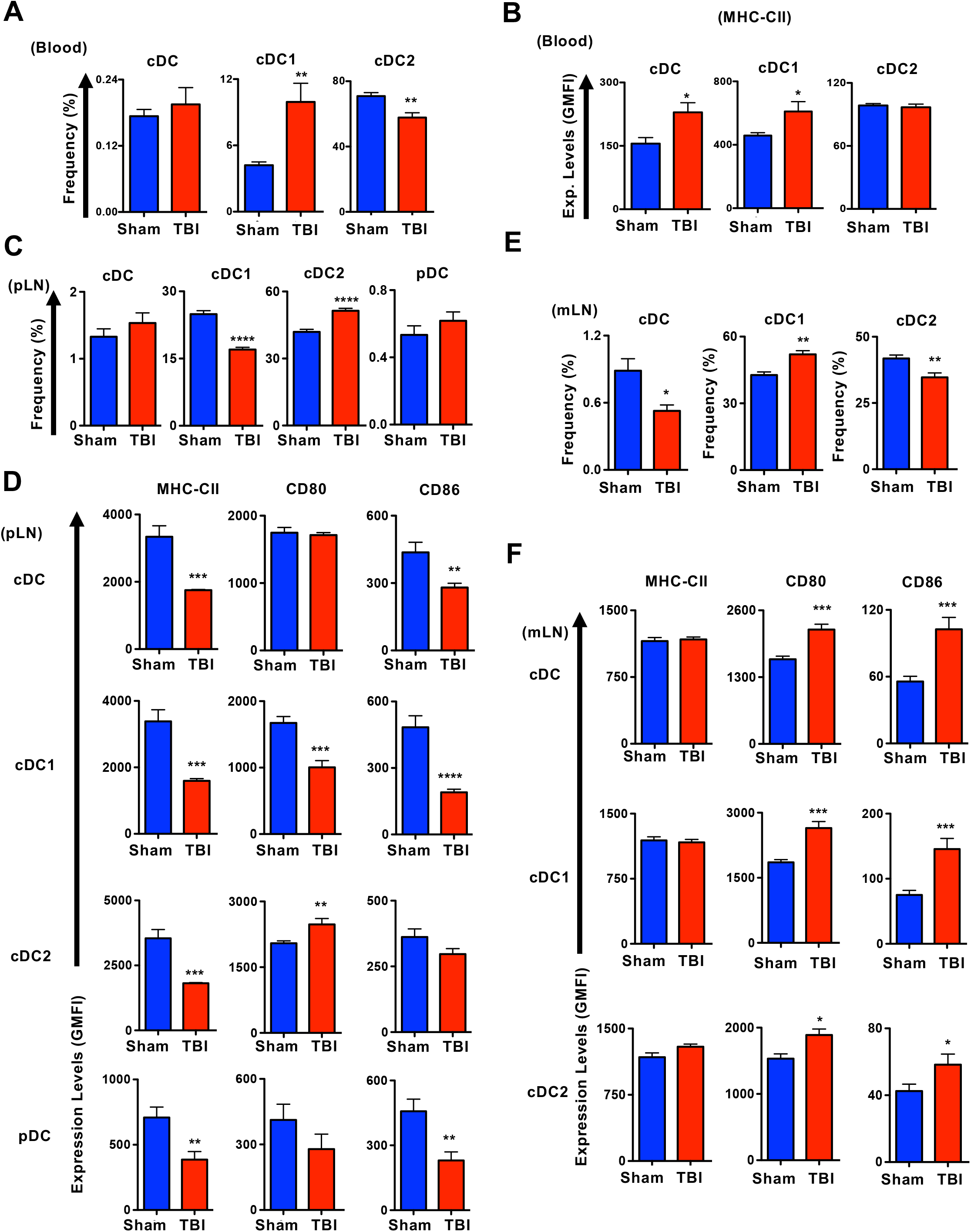
TBI results in an imbalance among DC subsets of the circulatory and lymphatic systems. **A**. Relative frequencies of total cDC, cDC1 and cDC2 subsets in the peripheral blood from Sham (n=4) and d7-TBI (n=5) mice. Data are representative of three independent experiments. **B**. Surface expression levels of MHC-CII in total cDC, cDC1 and cDC2 subsets of peripheral blood from Sham (n=4) and d7-TBI (n=5) mice. Shown were GMFI. Data are representative of three independent experiments. **C**. Relative frequencies of total cDC, cDC1, cDC2 pDC subsets in the peripheral lymph nodes from Sham (n=4) and d7-TBI (n=5) mice. Data are representative of two independent experiments. **D**. Surface expression levels of MHC-CII, CD80 and CD86 in total cDC, cDC1, cDC2 and pDC subsets of peripheral lymph nodes from Sham (n=4) and d7-TBI (n=5) mice. Shown were GMFI. Data are representative of two independent experiments. **E**. Relative frequencies of total cDC, cDC1 and cDC2 subsets in the mesenteric lymph nodes from Sham (n=4) and d7-TBI (n=5) mice. Data are representative of two independent experiments. **F**. Surface expression levels of MHC-CII, CD80 and CD86 in total cDC, cDC1, cDC2 and pDC subsets of mesenteric lymph nodes from Sham (n=4) and d7-TBI (n=5) mice. Shown were GMFI. Data are representative of two independent experiments. All data represent mean ± SEM. Two-tailed student’s t tests were used to assess statistical significance (*P < 0.05, **P<0.01, *** P< 0.001, **** P< 0.0001).

### TBI affects distribution of DC subsets within solid organs

DC subsets of the solid organs, including lungs, liver and skin, participate in active immune surveillance and maintaining a proper balance of cDC1, cDC2 and pDC subsets is vital for their functions[11, 17, 28, 29]. Emerging studies demonstrate that both non-lymphoid and lymphoid tissue DCs derive from the same precursors in the BM[11] and follow a related differentiation program[30, 31]. To test if TBI impacts the DC composition of solid organs, we determined the frequencies of DC compartments of the lungs. Frequencies of total CD11c^+^CII^+^ cDCs were increased within the CD45^+^ hematopoietic fraction of lungs from d7 TBI mice (**Fig. 5A**). However, within CD45^+^CD11c^+^CII^+^ cDCs, frequencies of CD103^+^ cDC1 subset were increased and CD11b^+^ cDC2 subset was normal in the lungs of d7 TBI mice (**Fig. 5B**). Further analysis of CD45^+^CD11c^+^CII^+^CD11b^+^ cDC2 fraction revealed a decrease of CD4^+^ cDC2 subset and normal frequencies of CD4^-^ cDC2 in the lungs of d7 TBI mice (**Fig. 5C**). Frequencies of CD45^+^CD11c^+^CII^+^ PDCA1^+^ pDCs were reduced in the lungs of d7 TBI mice (**Fig. 5D**).

**Figure 5.**
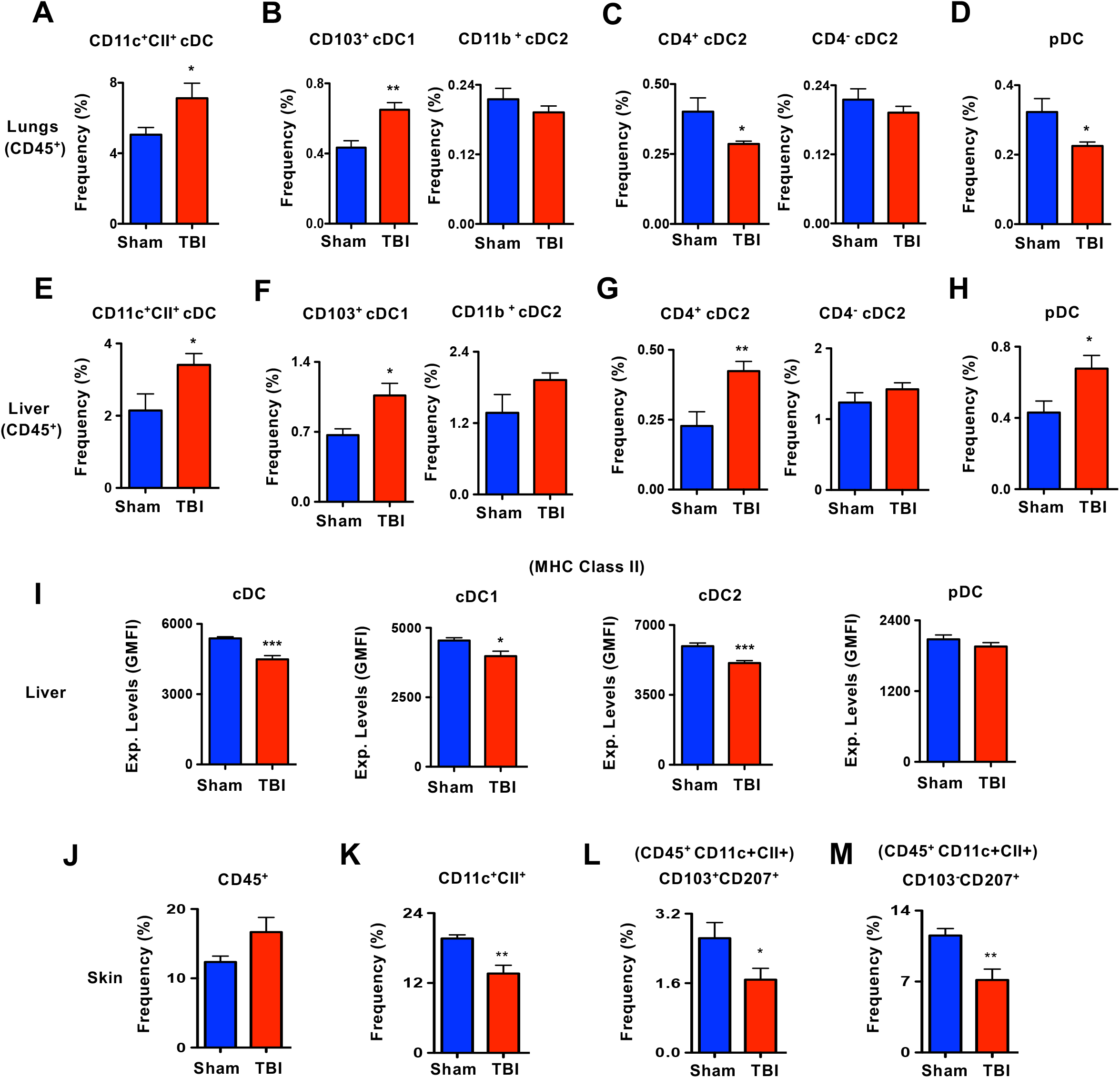
TBI affects DC distribution in solid organs. **A**. Frequencies of CD11c^+^CII^high^ cDCs within pre-gated CD45^+^ hematopoietic fraction of lungs from Sham (n=4) and d7-TBI (n=5) mice. Data are representative of two independent experiments. **B**. Frequencies of CD103^+^ cDC1 and CD11b^+^ cDC2 subsets within CD45^+^ CD11c^+^CII^high^ cDCs of lungs from Sham (n=4) and d7-TBI (n=5) mice. Data are representative of two independent experiments. **C**. Frequencies of CD4^+^ cDC2 and CD4^-^ cDC2 subsets within CD45^+^ CD11c^+^CII^high^ CD103^-^ CD11b^+^ cDC2 cells of lungs from Sham (n=4) and d7-TBI (n=5) mice. Data are representative of two independent experiments. **D**. Frequencies of CD11c^int^PDCA1^+^ pDCs within pre-gated CD45^+^ hematopoietic fraction of lungs from Sham (n=4) and d7-TBI (n=5) mice. Data are representative of two independent experiments. **E**. Frequencies of CD11c^+^CII^high^ cDCs within pre-gated CD45^+^ hematopoietic fraction of liver from Sham (n=4) and d7-TBI (n=5) mice. Data are representative of two independent experiments. **F**. Frequencies of CD103^+^ cDC1 and CD11b^+^ cDC2 subsets within CD45^+^ CD11c^+^CII^high^ cDCs of liver from Sham (n=4) and d7-TBI (n=5) mice. Data are representative of two independent experiments. **G**. Frequencies of CD4^+^ cDC2 and CD4^-^ cDC2 subsets within CD45^+^ CD11c^+^CII^high^ CD103^-^ CD11b^+^ cDC2 cells of liver from Sham (n=4) and d7-TBI (n=5) mice. Data are representative of two independent experiments. **H**. Frequencies of CD11c^int^PDCA1^+^ pDCs within pre-gated CD45^+^ hematopoietic fraction of liver from Sham (n=4) and d7-TBI (n=5) mice. Data are representative of two independent experiments. **I**. Surface expression levels of MHC-CII in total cDC, cDC1, cDC2 and pDC subsets of liver from Sham (n=4) and d7-TBI (n=5) mice. Shown were GMFI. Data are representative of two independent experiments. **J**. Frequencies of CD45^+^ hematopoietic cells from skin of Sham (n=4) and d7-TBI (n=5) mice. Data are representative of two independent experiments. **K**. Frequencies of CD11c^+^CII^high^ cDCs within pre-gated CD45^+^ hematopoietic fraction of skin from Sham (n=4) and d7-TBI (n=5) mice. Data are representative of two independent experiments. **L**. Frequencies of CD103^+^CD207^+^ dermal DCs within CD45^+^CD11c^+^CII^high^ cells of skin from Sham (n=4) and d7-TBI (n=5) mice. Data are representative of two independent experiments. **M**. Frequencies of CD103^-^CD207^+^ Langerhans Cells within CD45^+^CD11c^+^CII^high^ cells of skin from Sham (n=4) and d7-TBI (n=5) mice. Data are representative of two independent experiments. All data represent mean ± SEM. Two-tailed student’s t tests were used to assess statistical significance (*P < 0.05, **P<0.01, *** P< 0.001, **** P< 0.0001).

Next, we analyzed the DCs of liver from d7 TBI mice. Frequencies of CD11c^+^CII^+^ cDCs within CD45^+^ fraction were elevated in the liver of d7 TBI mice (**Fig. 5E**). However, within CD45^+^CD11c^+^CII^+^ cDCs, frequencies of CD103^+^ cDC1 subset were increased and CD11b^+^ cDC2 subset was normal in the liver of d7 TBI mice (**Fig. 5F**). Further analysis of CD45^+^CD11c^+^CII^+^CD11b^+^ cDC2 fraction revealed an increase of CD4^+^ cDC2 subset and normal frequencies of CD4^-^ cDC2 in the liver of d7 TBI mice (**Fig. 5G**). Frequencies of CD45^+^CD11c^+^CII^+^ PDCA1^+^ pDCs were elevated in the liver of d7 TBI mice (**Fig. 5H**). Immunophenotyping studies indicated reduced expression levels of MHC-CII in total cDC, cDC1 and cDC2 subsets, whereas normal expression levels of MHC-CII in pDCs from the liver of d7 TBI mice (**Fig. 5I**).

Finally, we determined the frequencies of DCs in the skin. Frequencies of total CD45^+^ hematopoietic cells were increased in the skin of d7 TBI mice (**Fig. 5J**). However, within CD45^+^ fraction, the frequencies of CD11c^+^CII^+^ DCs were reduced in the skin of d7 TBI mice (**Fig. 5K**). Determination of CD11c^+^CII^+^ CD103^+^CD207^+^ dermal DCs and CD11c^+^CII^+^ CD103^-^ CD207^+^ Langerhans Cells (LCs) indicated a reduction of both fractions in the skin of d7 TBI mice (**Fig. 5L-M**). Overall, these data indicate that TBI impacts the distribution and composition of DC subsets within the solid organs that participate in immune-surveillance functions.

### TBI affects early differentiation pathways of DCs in the BM

Early stages of DC differentiation occur in the BM through commitment of hematopoietic stem cells (HSCs) to common dendritic cell progenitors (CDPs)[32]. Indeed, changes in the frequencies and numbers of CDPs result in abnormal composition of DCs in both lymphoid and non-lymphoid organs[11, 16, 17]. In view of the fact that TBI affects distribution of DCs in almost all organs, we investigated if TBI has an impact on the early stages of DC differentiation. Flow cytometric analyses identified reduced relative frequencies of Lin^-^c-Kit^int^Flt3^+^CD115^+^ CDPs in the BM of d1 and d3 TBI mice (**Fig. 6A-B**), decreased overall frequencies of CDPs in the BM of d3 TBI mice (**Fig. 6A,C**), and diminished absolute numbers of CDPs in the BM of d3 TBI mice (**Fig. 6A,D**). On the other hand, relative frequencies (**Fig. 6A, E**), overall frequencies (**Fig. 6A, F**), and absolute numbers (**Fig. 6A, G**) of Lin^-^c-Kit^int^Flt3^+^CD115^-^ pDC-committed-progenitors were augmented in the BM of both d1 and d3 TBI mice. To study if these changes in the DC progenitors of the BM are consistent at the later phase of TBI, we analyzed the BM of d7 TBI mice. Interestingly, our data on CDPs indicated that the relative frequencies were normal (**Fig. 6H**), overall frequencies were increased (**Fig. 6I**), and the absolute numbers (**Fig. 6J**) were augmented in the BM of d7 TBI mice. However, the relative frequencies (**Fig. 6K**), overall frequencies (**Fig. 6L**), and absolute numbers (**Fig. 6M**) remain normal in the BM of d7 TBI mice. These data demonstrate that TBI affects early differentiation program of DCs, especially at the stages of commitment to CDPs.

**Figure 6.**
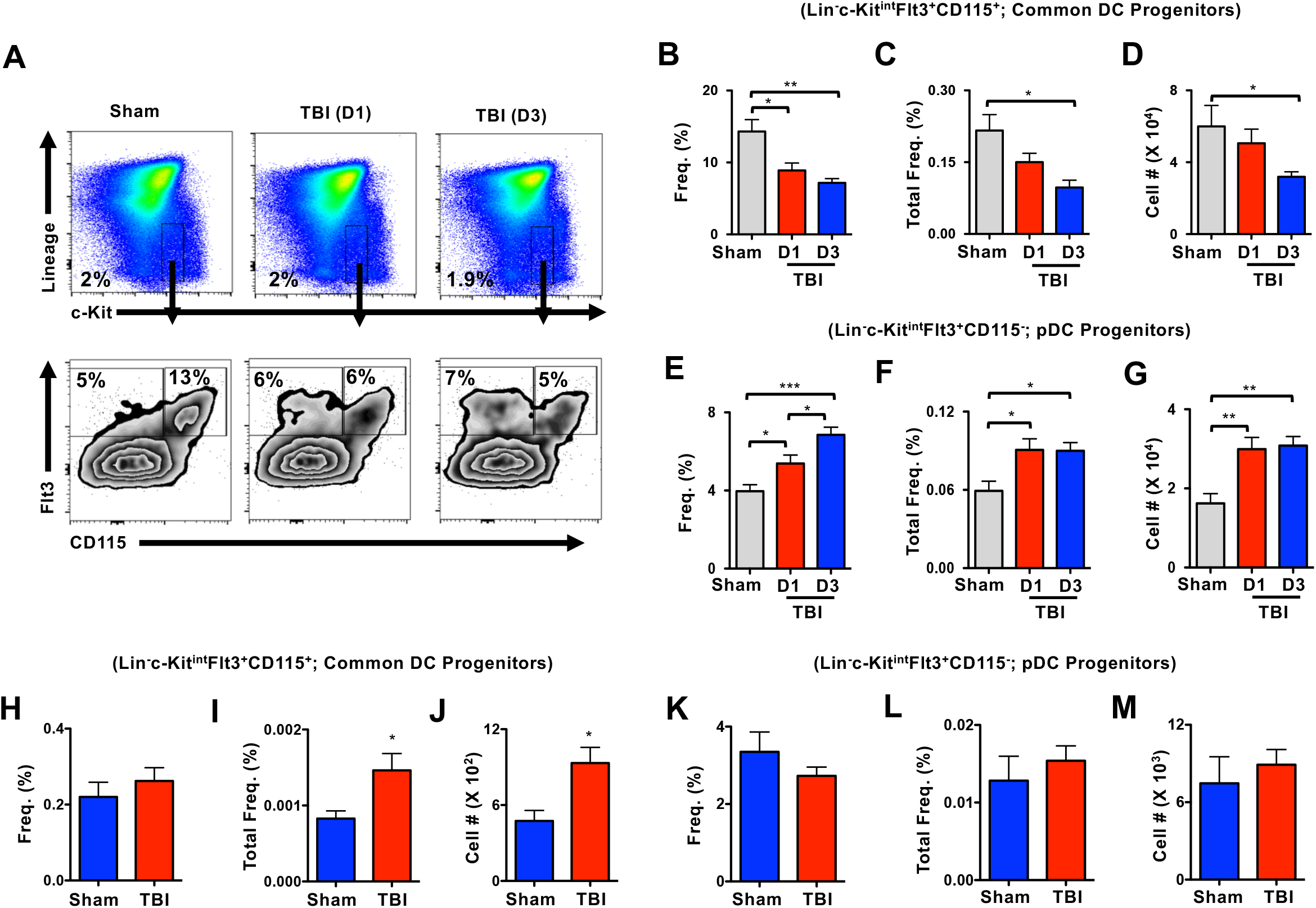
TBI affects early differentiation of DC progenitors in the BM. **A**. FACS plots indicating frequencies of Lin^-^c-Kit^+^ Flt3^+^CD115^+^ CDPs and Lin^-^c-Kit^+^ Flt3^+^CD115^-^ pDC progenitors in the BM of Sham (n=4), d1-TBI (n=5) and d3-TBI (n=5) mice. Data are representative of two independent experiments. **B-D**. Relative frequencies (**B**), overall frequencies (**C**) and absolute numbers (**D**) of Lin^-^c-Kit^+^ Flt3^+^CD115^+^ CDPs in the BM of Sham (n=4), d1-TBI (n=5) and d3-TBI (n=5) mice. Data are representative of two independent experiments. **E-G**. Relative frequencies (**E**), overall frequencies (**F**) and absolute numbers (**G**) of Lin^-^c-Kit^+^ Flt3^+^CD115^-^ pDC progenitors in the BM of Sham (n=4), d1-TBI (n=5) and d3-TBI (n=5) mice. Data are representative of two independent experiments. **H-J**. Relative frequencies (**H**), overall frequencies (**I**) and absolute numbers (**J**) of CDPs in the BM of Sham (n=4) and d7-TBI (n=5) mice. Data are representative of two independent experiments. **K-M**. Relative frequencies (**K**), overall frequencies (**L**) and absolute numbers (**M**) of pDC progenitors in the BM of Sham (n=4) and d7-TBI (n=5) mice. Data are representative of two independent experiments. All data represent mean ± SEM. Mann-Whitney non-parametric tests and two-tailed student’s t tests were used to assess statistical significance (*P < 0.05, **P<0.01, *** P< 0.001, **** P< 0.0001).

### TBI suppresses Reactive Oxygen Species production in DC-subsets and progenitors

One of the hallmark features of secondary injury following TBI is augmented levels of ROS and oxidative stress in innate immune cells, including microglia and macrophages[33, 34]. Of note, ROS have been shown to play critical and decisive roles at the early differentiation stages of human and mouse DCs in the BM[35-39]. In an attempt to understand the molecular mechanisms that could contribute to altered DC differentiation at early and late phases of TBI, we determined the expression levels of ROS in DC subsets and progenitors. Intriguingly, our analysis indicated reduced frequencies of ROS^high^ and total intracellular ROS levels in cDCs of spleen on d1 and d3 after TBI (**Fig. 7A, B**). Further analysis of cDC subsets revealed a reduction in the frequency of ROS^high^ splenic cDC1 subset from d1 TBI mice (**Fig. 7C, D**), and a decrease in both frequencies of ROS^high^ and total intracellular ROS levels in splenic cDC2 subset from d1 and d3 TBI mice (**Fig. 7E, F**). Consistently, analysis of CD8^-^CD11b^-^ immature cDCs exhibited reduced frequencies of ROS^high^ cells and total intracellular ROS levels from d1 and d3 TBI mice (**Fig. 7G, H**). On the other hand, ROS levels in pDCs indicated normal frequencies of ROS^high^ pDCs from d1 and d3 TBI mice and a reduction in total intracellular ROS levels in pDCs from d3 TBI mice (**Fig. 7I, J**). To assess the alterations in ROS levels in DC subsets at late phase of TBI, we studied DCs from d7 TBI mice. Our data concluded that the total intracellular ROS levels were remarkably reduced in total cDC (**Fig. 7K**), cDC1 (**Fig. 7L**), cDC2 (**Fig. 7M**), immature cDC (**Fig. 7N**) and pDC (**Fig. 7O**) subsets from the spleen of d7 TBI mice. Finally, we determined if ROS levels are altered at the early development stages of DCs following TBI. Analysis indicated that both frequencies of ROS^high^ CDPs and total intracellular ROS levels in CDPs were reduced in the BM of d1 and d3 TBI mice (**Fig. 7P**). In essence, these data established that the intracellular ROS levels were reduced in DC progenitors and differentiated DCs at both early and late phases of TBI.

**Figure 7.**
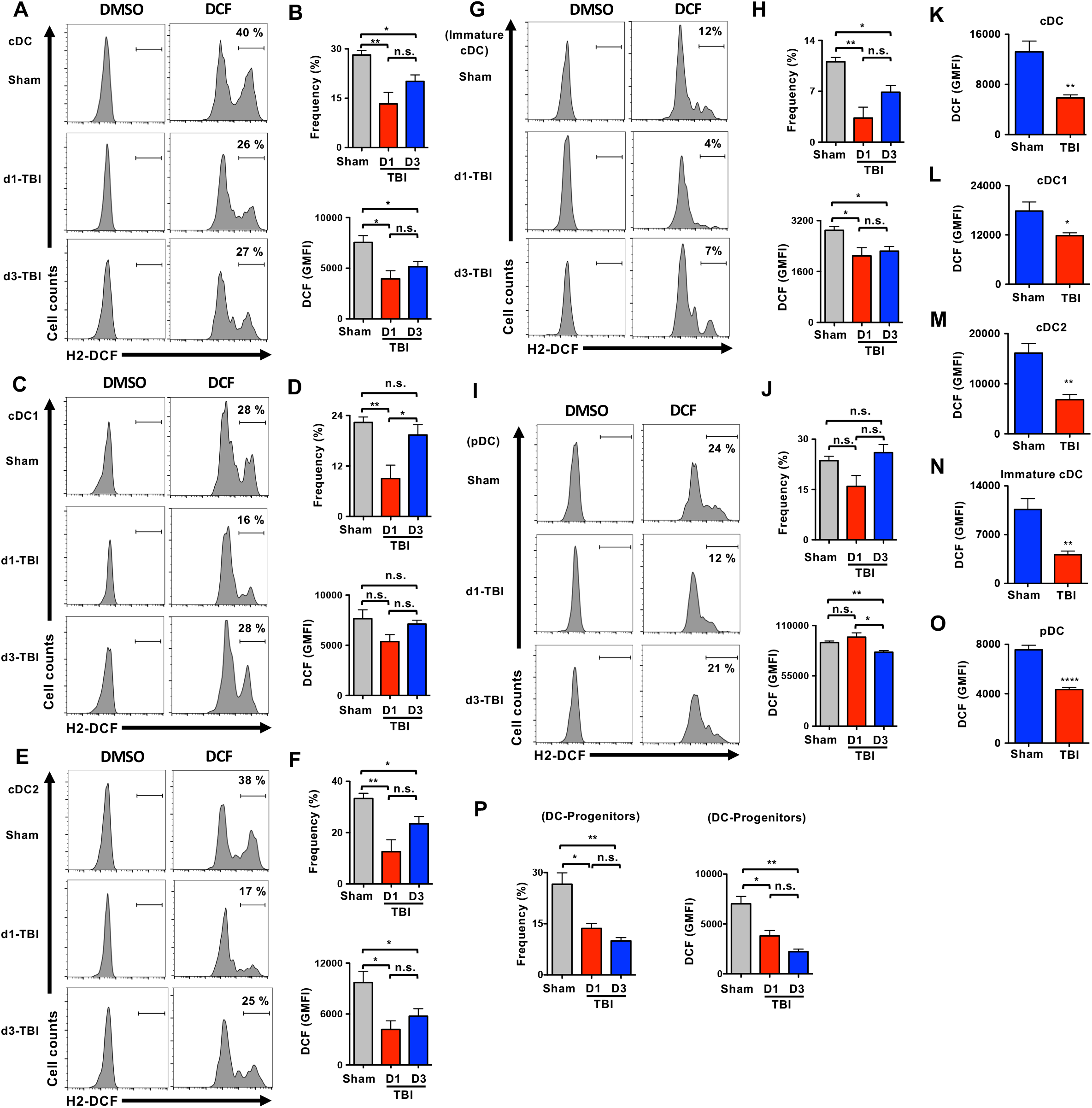
TBI results in diminished levels of ROS in DC subsets and Progenitors. **A**. Histograms indicating intracellular ROS levels in CD11c^+^CII^+^ splenic cDCs from sham (n=4), d1-TBI (n=5) and d3-TBI (n=5) mice. Shown were the frequencies of ROS^high^ cells under the indicated gates. Cells treated with DMSO served as negative controls. **B**. Frequencies of gated ROS^high^ CD11c^+^CII^+^ splenic cDCs (**top**) and total intracellular ROS levels (GMFI) in CD11c^+^CII^+^ splenic cDCs (**bottom**) from sham (n=4), d1-TBI (n=5) and d3-TBI (n=5) mice. **C**. Histograms indicating intracellular ROS levels in CD8^+^CD11b^-^CD11c^+^CII^+^ splenic cDC1 subset from sham (n=4), d1-TBI (n=5) and d3-TBI (n=5) mice. Shown were the frequencies of ROS^high^ cells under the indicated gates. Cells treated with DMSO served as negative controls. **D**. Frequencies of gated ROS^high^ CD8^+^CD11b^-^CD11c^+^CII^+^ splenic cDC1 subset (**top**) and total intracellular ROS levels (GMFI) in CD8^+^CD11b^-^CD11c^+^CII^+^ splenic cDC1 subset (**bottom**) from sham (n=4), d1-TBI (n=5) and d3-TBI (n=5) mice. **E**. Histograms indicating intracellular ROS levels in CD8^-^CD11b^+^CD11c^+^CII^+^ splenic cDC2 subset from sham (n=4), d1-TBI (n=5) and d3-TBI (n=5) mice. Shown were the frequencies of ROS^high^ cells under the indicated gates. Cells treated with DMSO served as negative controls. **F**. Frequencies of gated ROS^high^ CD8^-^CD11b^+^CD11c^+^CII^+^ splenic cDC2 (**top**) and total intracellular ROS levels (GMFI) in CD8^-^CD11b^+^CD11c^+^CII^+^ splenic cDC2 (**bottom**) from sham (n=4), d1-TBI (n=5) and d3-TBI (n=5) mice. **G**. Histograms indicating intracellular ROS levels in CD8^-^CD11b^-^CD11c^+^CII^+^ splenic immature cDCs from sham (n=4), d1-TBI (n=5) and d3-TBI (n=5) mice. Shown were the frequencies of ROS^high^ cells under the indicated gates. Cells treated with DMSO served as negative controls. **H**. Frequencies of gated ROS^high^ CD8^-^CD11b^-^CD11c^+^CII^+^ splenic immature cDCs (**top**) and total intracellular ROS levels (GMFI) in CD8^-^CD11b^-^CD11c^+^CII^+^ splenic immature cDCs (**bottom**) from sham (n=4), d1-TBI (n=5) and d3-TBI (n=5) mice. **I**. Histograms indicating intracellular ROS levels in PDCA1^+^CD11c^+^ splenic pDCs from sham (n=4), d1-TBI (n=5) and d3-TBI (n=5) mice. Shown were the frequencies of ROS^high^ cells under the indicated gates. Cells treated with DMSO served as negative controls. **J**. Frequencies of gated ROS^high^ PDCA1^+^CD11c^+^ splenic pDCs (**top**) and total intracellular ROS levels (GMFI) in PDCA1^+^CD11c^+^ splenic pDCs (**bottom**) from sham (n=4), d1-TBI (n=5) and d3-TBI (n=5) mice. **K-O**. Histograms indicating intracellular ROS levels (GMFI) in total cDCs (**K**), cDC1 (**L**), cDC2 (**M**), immature cDCs (**N**) and pDCs (**O**) of the spleen from sham (n=5) and d7-TBI (n=5) mice. **P**. Frequencies of gated ROS^high^ Lin^-^c-Kit^+^ Flt3^+^CD115^+^ CDPs (**left**) and total intracellular ROS levels (GMFI) in Lin^-^c-Kit^+^ Flt3^+^CD115^+^ CDPs (**right**) of the BM of Sham (n=4), d1-TBI (n=5) and d3-TBI (n=5) mice. Data are representative of two independent experiments. All data represent mean ± SEM. Mann-Whitney non-parametric tests and two-tailed student’s t tests were used to assess statistical significance (*P < 0.05, **P<0.01, *** P< 0.001, **** P< 0.0001)

## Discussion

Infections are a leading cause of morbidity and mortality in patients with acute CNS injury, such as TBI, stroke and spinal cord injury[19-21, 40]. Indeed, there exists a direct correlation between infections and poorer clinical outcomes overall[41]. More importantly, infections were found to be the key driver of ∼ 19% of mortality in TBI patients, that survived at least one year after TBI[42]. Mounting evidences indicate that nosocomial infections (following TBI) are major contributors of not only to mortality, but also inducers of deleterious long-term consequences in patients[43]. Furthermore, infections acquired after TBI may trigger pathophysiological sequelae and exacerbate neurodegeneration[44, 45], intracranial hypertension and an associated need for longer duration on mechanical ventilation[46-50]. Indeed, survivors of brain trauma are 2.3-4.3 times more likely to die than general population, with increased risk of death from pneumonia or sepsis[42]. Recent studies indicated that up to 75% of severe TBI patients have been reported to develop sepsis, that itself is linked to high rates of morbidity, cognitive impairment, anxiety and depression, mediated multi-organ failure[51]. Post-traumatic immune insults are life-threatening complications in patients with moderate-severe TBI and observed in up to 55 % of patients[51]. The most common nosocomial infections in TBI patients include, pneumonia/lower respiratory tract infections, urinary tract infections and surgical site infections[19, 20, 51-53]. Given the fact that resistance to all of these infections is dependent on a functional immune system, providing and/or preserving competent and functional immune components would be a reasonable and effective therapeutic strategy to treat TBI patients.

DCs are regarded as the most professional antigen presenting cells and indispensable in initiating T-, B- and NK-cell mediated adaptive immune responses against viral, fungal and bacterial infections[9-12]. In particular, DCs are essential to provide immunity against lung infections, including viral and bacterial pneumonia[28, 54-58]. Intriguingly, our data from this current study document that TBI directly affects the pool and composition of DCs in almost all lymphoid and non-lymphoid organs, including lungs. These altered frequencies and numbers of cDC1, cDC2 and pDCs subsets in each tissue may be, at least in part, responsible for the immune dysfunction caused by TBI. More importantly, our analyses concluded that TBI affects the immunogenicity of DCs, such as altered surface expression of MHC CI and CII proteins and co-stimulatory molecules-CD80 and CD86, which may eventually lead to defective functions of DCs.

Mounting evidences established that TBI affects organs outside the CNS. Mirzayan *et al*. reported that TBI causes augmented migration and infiltration of immune cells to peripheral organs, such as lungs and liver, which eventually led to histopathological changes and various degrees of organ dysfunction[59]. Interestingly, their study concluded that the composition of spleen and kidney were not altered in response to TBI. On the other hand, a series of studies established that peripheral immune cells play a critical role in the overall pathology after brain injury. Indeed, splenectomy in rats immediately after TBI resulted in decreased expression of pro-inflammatory cytokines, mortality rate, improved cognitive function[60] and attenuated neurodegeneration and CCL20 chemokine expression in the brain[18]. Most of the peripheral innate immune cells, including neutrophils, monocytes and macrophages, are believed to respond at the early phase and decline within few days after TBI[61]. Intriguingly, our present data suggest that peripheral DC composition and numbers are altered within 24 hours of TBI and exhibited sustained changes even after 7 days of TBI. However, it is unclear, at present, if peripheral DCs migrate to the injured site in the brain and participate in inflammation and neurodegeneration. Future studies with an in-depth focus on identifying the involvement of DCs in neuroinflammation and pathophysiology of CNS would be essential.

Emerging studies have unequivocally established the contribution of oxidative stress to secondary injury in TBI pathology, including development of cerebral edema, inflammation, and the secondary neuronal damage found post-TBI. Disrupted blood flow after TBI leads to cerebral hypoxia or ischemia with the consequent decrease of oxygen and glucose supply to the CNS[62]. In addition, mitochondrial dysfunction caused by TBI results in marked accumulation of ROS in the brain[63]. In sharp contrast, our studies identified that ROS levels were rather diminished in both differentiated DCs and DC subsets at early and late phases of TBI. In keeping with our findings, neutrophil ROS production in patients with moderate or severe TBI was significantly lower than that of healthy age- and sex-matched individuals[61]. Given the importance of ROS in the differentiation and functional integrity of DCs[35-39], we speculate that impaired ROS generation might be responsible for DC alterations in peripheral organs after TBI. At present, it is unclear as if TBI is either directly or indirectly responsible for the ROS defects in DCs. In addition, mechanisms that are responsible for reduced ROS in DCs after TBI need to be elucidated.

While a great deal of research has been devoted to deciphering the relationship between brain trauma and peripheral immune system, very little is known regarding precise mechanisms through which interactions between peripheral lymphoid organs and CNS are regulated. Neuronal and hormonal mechanisms, such as the cholinergic anti-inflammatory pathway, the adrenergic pathway and the hypothalamic pituitary axis (HPA), have been postulated to play crucial roles in regulating the peripheral immune response following CNS injury[20, 64-66]. Interestingly, most of these studies have focused on brain-spleen crosstalk following CNS injury[65, 67]. It is unclear if such neuronal and hormonal mechanisms play any roles in the early differentiation pathways of immune cells in BM following TBI. In the present study, we establish that the alterations of DCs in the peripheral lymphoid organs, including spleen, are associated with increased differentiation of CDPs in the BM. Of note, TBI induced inflammatory response is not confined to the brain and can cause a systemic inflammatory response syndrome (SIRS), including elevated levels of TNFα, IL1β and IL6 in the blood serum[68-71]. Recent studies, including our own[24-27, 72], demonstrated that exaggerated expression of inflammatory cytokines, such as TNFα, IL1β, IFNα, IFNγ, and IL6, affects early differentiation of immune cells[73-75]. Based on these studies, it is tempting to speculate that exaggerated pro-inflammatory signals might be responsible, at least in part, for increased early differentiation of DCs in the BM after TBI. However, the involvement of additional mechanisms, including neuronal and hormonal control, in the differentiation and maintenance DCs after CNS injury cannot be underestimated and awaits further investigation. A deeper understanding on the molecular mechanisms that contribute to DC defects following TBI would be essential and beneficial in treating infections in patients with acute central nervous system (CNS) injuries, such as TBI, stroke and spinal cord injury.

## Conclusion

Our data demonstrate, for the first time, that TBI affects the distribution pattern of DCs and induces an imbalance among DC subsets in both lymphoid and non-lymphoid organs. In addition, the current study demonstrates that TBI results in reduced levels of ROS in DCs at both early and late phases of TBI, which may explain altered DC differentiation paradigm following TBI. A deeper understanding on the molecular mechanisms that contribute to DC defects following TBI would be essential and beneficial in treating infections in patients with acute CNS injuries, such as TBI, stroke and spinal cord injury.

## Availability of data and materials

The data that support the findings of this study are available from corresponding author on reasonable request.

## Abbreviations

TBI: Traumatic Brain Injury
DC: Dendritic Cells
CDP: Common Dendritic Cell Progenitor
pDC-P: plasmacytoid DC Progenitor
cDCs: conventional or classical DCs
pDCs: plasmacytoid DCs
CNS: Central Nervous System
ROS: Reactive Oxygen Species
BM: Bone Marrow
pLN: peripheral lymph node
mLN: peripheral lymph node

## Consent to participate

Not applicable

## Acknowledgements

Not applicable

## Contributions

O.T. performed animal surgery; V.G. helped with animal surgery; J.M.S. provided help with animal surgery; C.V.R. designed and performed research, collected all data, analyzed and interpreted data, prepared all figures and wrote the manuscript. All authors reviewed the manuscript.

## Ethics declarations

### Ethics approval

This study was approved by the University of Maryland School of Medicine Institutional Animal Care and Use Committee (IACUC). All experiments were performed in accordance with the National Institutes of Health Guidelines on the Use of Laboratory Animals.

### Consent for publication

Not applicable

